# Reciprocal nutritional benefits in a sponge-seagrass association

**DOI:** 10.1101/2024.12.06.627200

**Authors:** U. Cardini, L.M. Montilla, G. Zapata-Hernández, J. Berlinghof, E. Guarcini, M. Furia, F. Margiotta, T.B. Meador, C. Wild, S. Fraschetti, I. Olivé

## Abstract

Sponges commonly form associations within seagrass meadows, but their potential impact on seagrass productivity and nutrient cycles remains poorly understood. This study investigates the association between the demosponge *Chondrilla nucula* and the Mediterranean seagrass *Posidonia oceanica* in two sampling occasions during the plant growth (spring) and senescence (autumn) seasons at a small inlet near Naples, Italy, where the sponge grows conspicuously within the seagrass bed. We found a non-linear relationship between the benthic cover of the sponge and the seagrass, with higher *C. nucula* cover linked to intermediate *P. oceanica* cover, suggesting spatial dependence. *P. oceanica* showed higher net primary production (NPP) in spring, while *C. nucula* was net heterotrophic in spring but exhibited slightly positive NPP in autumn. NPP remained stable when the two organisms were associated, regardless of the season. *C. nucula* consistently contributed inorganic nutrients to the association in the form of phosphate, ammonium, and substantial nitrate, recycling nutrients that potentially benefited *P. oceanica* in its growth season. In return, the seagrass consistently provided dissolved organic carbon, which aided sponge nutrition in spring. These findings suggest reciprocal benefits in the interaction between *C. nucula* and *P. oceanica*, with nutrient exchange facilitating a facultative mutualism that potentially supports and stabilizes the productivity of the seagrass ecosystem.

**SIGNIFICANCE STATEMENT:** This study provides a novel exploration of the reciprocal interactions between the demosponge *Chondrilla nucula* and the Mediterranean seagrass *Posidonia oceanica*, revealing a facultative mutualism mediated by nutrient exchange. Our findings show a non-linear spatial dependence between sponge and seagrass cover and demonstrate the sponge’s substantial contributions of inorganic nutrients (phosphate, ammonium and conspicuous nitrate) to the seagrass, particularly during its productive spring season. In return, *P. oceanica* supplies dissolved organic matter, aiding sponge nutrition. This study uniquely quantifies these reciprocal nutrient exchanges across the plant growth and senescence seasons, demonstrating how such interactions stabilize net primary production and support ecosystem functioning. These insights address a critical gap in understanding the role of sponge-seagrass associations in nutrient cycling, highlighting their significance for the resilience, productivity, and metabolic balance of coastal ecosystems under changing environmental conditions.

## INTRODUCTION

Seagrasses are vital ecosystem engineers that create habitats for diverse marine life. Seagrass meadows support significantly more species than unvegetated areas, particularly among fish and invertebrate communities [1], while fostering complex epibenthic assemblages [2]. Many invertebrates use these meadows as sources of organic matter [3] and as shelter [4], and they also host diverse microbiomes that perform key ecosystem functions, contributing to so-called nested ecosystems [5-7]. Positive species interactions are recognized as crucial drivers of community structure and ecosystem functioning in seagrass ecosystems [7-10]. However, although this topic has been well explored in terrestrial environments, substantial knowledge gaps remain in marine systems [e.g., 11].

Among seagrass-associated invertebrates, sponges play a crucial role in nutrient cycling. They consume dissolved organic carbon (DOC) and transform it into detritus, making it available for higher trophic levels through what is known as the sponge loop [12, 13]. High-microbial abundance (HMA) sponges, in particular, absorb dissolved organic matter (DOM) and release nitrate 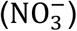, ammonium 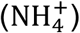, and phosphate 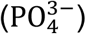, which are essential for nutrient recycling [14]. This function makes sponges beneficial partners for primary producers (PP), such as seagrasses and macroalgae, which release large amounts of DOM into their surroundings [15]. At the same time, the growth of PP is often limited by inorganic nitrogen [16], underscoring the potential significance of sponge-PP interactions.

Sponge-PP associations have been documented in various marine environments. On coral reefs, sponges absorb DOC from corals and macroalgae, returning inorganic nutrients in a reciprocal, mutually beneficial relationship [13, 17, 18]. Similar associations occur in mangrove ecosystems, where sponges release nitrogen that supports mangrove growth, while receiving carbon from mangrove roots, establishing facultative mutualisms [19].

In seagrass meadows, sponges have been shown to enhance growth and nutrient content of primary producers, as demonstrated in the association of the sponge *Ircina felix* with the seagrass *Thalassia testudinum* and other non-dominant seagrass species [20]. Another example involves *T. testudinum* benefiting from nutrients like ammonium 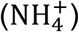 and phosphate 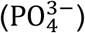 released by the sponge *Halichondria melanadocia* [21, 22]. However, despite growing research interest in sponge-PP associations, only a limited number of studies have been dedicated to these interactions to date, leaving key ecological processes insufficiently understood. In particular, a quantification of the effect of sponge-seagrass associations on seagrass productivity and nutrient cycles remains largely unexplored, and the extent to which these interactions enhance or stabilize seagrass primary production under varying environmental conditions is still unclear. Addressing these gaps is essential for understanding the resilience of seagrass ecosystems and their capacity to adapt to global change.

In the Mediterranean Sea, the seagrass *Posidonia oceanica* forms extensive meadows that extend from the surface down to about 40 m depth. Among the variety of associated biodiversity, shallow *P. oceanica* meadows frequently host the demosponge *Chondrilla nucula* [23]. *C. nucula* is widespread across the Mediterranean Sea and is classified as a high-microbial-abundance (HMA) sponge [24], harboring a rich and diverse microbiome dominated by Cyanobacteria [25]. Similar microbiome compositions have been found in *C. nucula* populations across other Mediterranean locations [26] and in the congeneric *C. caribensis* from the Caribbean [27], suggesting stable core bacterial assemblages in *Chondrilla* spp. across regions. Phylogenetic analyses identified cyanobacterial symbionts in *C. nucula* and proposed them as *Candidatus Synechococcus spongiarum* for *C. nucula* from the Mediterranean and *C. australiensis* from Australia [28]. In the Caribbean species *C. caribensis*, these symbionts provide nutritional benefits through photosynthate translocation and algal cell ingestion [29].

This study investigates the association between *C. nucula* and *P. oceanica* in the central Tyrrhenian Sea (Italy), focusing on a coastal site in the Gulf of Pozzuoli. Here, *C. nucula* grows abundantly at the base of seagrass shoots and expands laterally to adjacent rhizomes, despite the availability of alternative substrates nearby (Fig. 1). We explore whether the *C. nucula-P. oceanica* association can be characterized as a facultative mutualism, hypothesizing that DOM released by *P. oceanica* supports sponge nutrition, while the sponge provides a source of inorganic nutrients to the seagrass.

**Figure 1.**
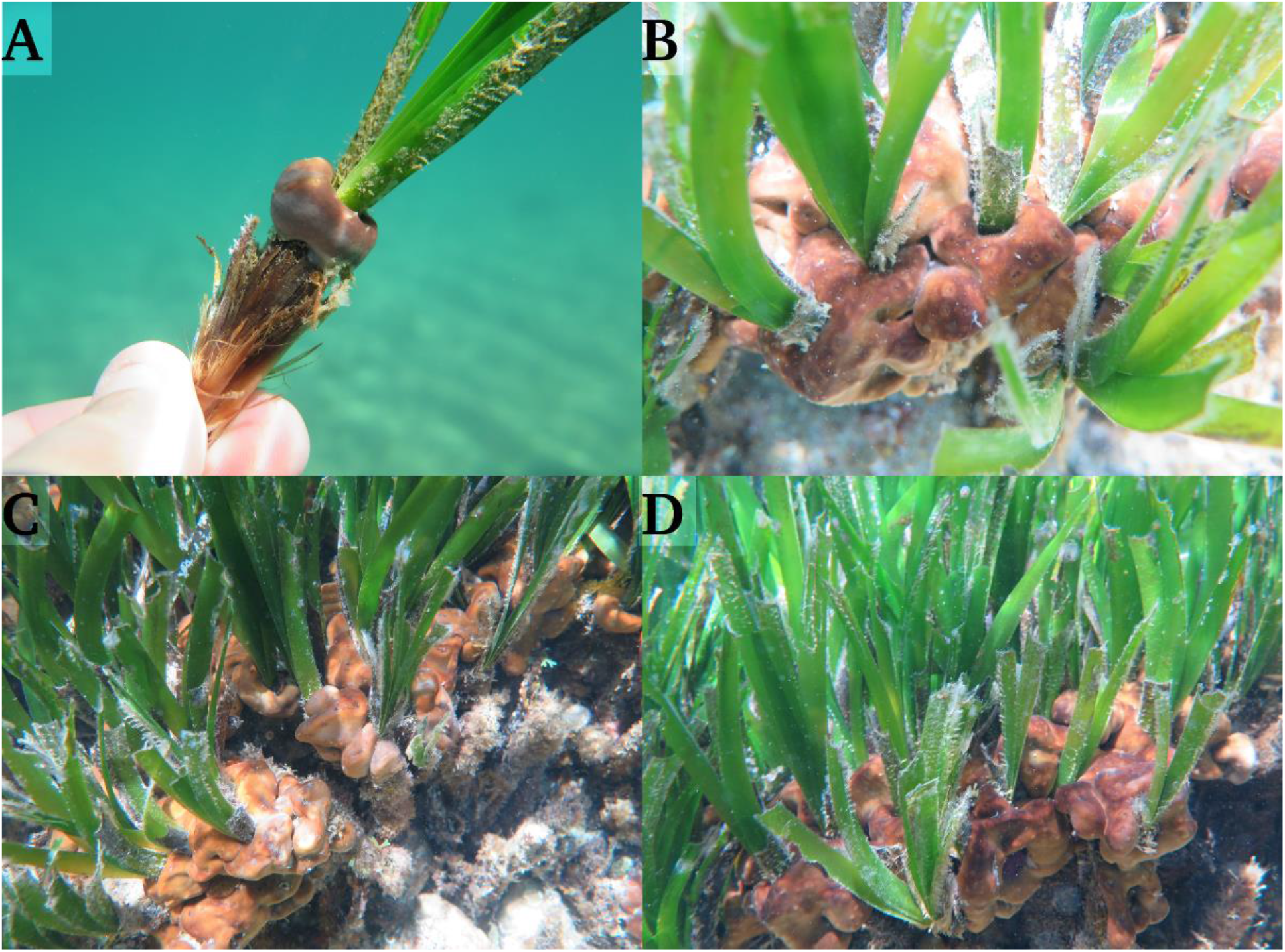
Photographic documentation of the association between *Posidonia oceanica* and *Chondrilla nucula* at the study site. (A) Close-up of a *P. oceanica* shoot with a *C. nucula* bundle surrounding the transition zone between the leaves and foliar sheath. (B) Detail of sponge bundles firmly attached to individual shoots. (C, D) Large *C. nucula* colonies covering multiple shoots and fusing into extensive, contiguous growths. Photos: U. Cardini.

To test this hypothesis, we: (i) conducted a spatial distribution analysis of *C. nucula* within the seagrass meadow, (ii) quantified net fluxes of oxygen, organic, and inorganic nutrients in closed chamber incubations, and (iii) used stable isotope analyses to examine potential signals of nutrient transfer in the sponge-seagrass association. These experiments evaluated the effect of each organism, both individually and in association, during the plant growth (spring) and senescence (autumn) seasons. Together, these approaches helped to gain clarity on the nature of this sponge-seagrass association and determine whether nutrient exchange plays a role, supporting the characterization of this association as a facultative mutualism.

## RESULTS

### Environmental conditions and seagrass-sponge association patterns

Environmental variables at the sampling site were higher in autumn than in spring for 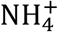, nitrate+nitrite 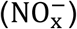, and DOC concentrations, as well as for seawater temperature (Table 1). The relationship between seagrass and sponge cover was non-linear (Fig. 2), and our model revealed a cubic curve (edf = 3.7, p = 6.55 × 10^−5^), with a peak in sponge cover (∼10% and up to 30%) at intermediate levels of seagrass cover (∼75%; Fig. 2). The asymmetry analysis supported this trend with significant q coefficients (Table S1).

**Table 1.** Environmental conditions at the study site in the two seasons at the time of the incubation experiments. Bold *p* values indicate significant differences between the two seasons in the respective variable (t-test).

| Season | NH<sub>4</sub><sup>+</sup> (μM) | NO<sub>x</sub><sup>-</sup> (μM) | PO<sub>4</sub><sup>3-</sup> (μM) | DOC (μM) | DON (μM) | Light intensity (LUX) | Temperature (°C) |
| --- | --- | --- | --- | --- | --- | --- | --- |
| Autumn | 0.98 ± 0.01 | 0.97 ± 0.08 | 0.05 ± 0.01 | 117.3 ± 12.5 | 6.4 ± 2.9 | 49371 ± 42228 | 19.5 ± 0.1 |
| Spring | 0.39 ± 0.05 | 0.66 ± 0.03 | 0.06 ± 0.01 | 81.9 ± 4.0 | 6.2 ± 0.2 | 66313 ± 34418 | 20.2 ± 0.1 |
| *p* value | **0.003** | **0.009** | 0.129 | **0.004** | 0.943 | 0.314 | **<0.001** |

**Figure 2.**
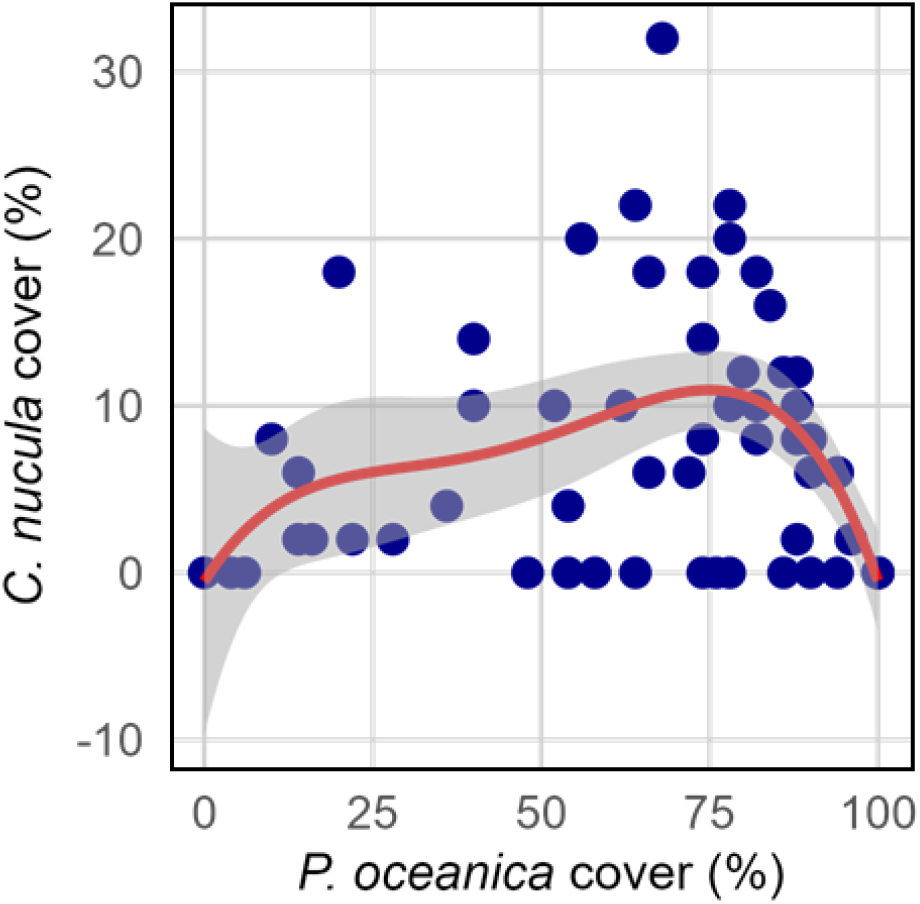
Relationship between *P. oceanica* cover (%) and *C. nucula* cover (%) based on field observations. Each point represents a discrete measurement, with *P. oceanica* cover plotted on the x-axis and *C. nucula* cover on the y-axis. A fourth-degree polynomial regression (red line) models the relationship, with shaded gray ribbons representing the 95% confidence interval.

### Primary production and respiration rates

Hourly rates of net primary production (NPP) were highest in the seagrass, followed by the seagrass-sponge association, and lowest in the sponge (Fig. 3A, Table S2). A seasonal effect was particularly evident in the seagrass, where NPP in autumn was 37% lower than in spring (11.1 ± 3.6 vs 17.5 ± 3.5 µmol O_2_ g DW−^1^ h−^1^, respectively; Fig. 3A, Table S3). Conversely, NPP rates in the sponge were higher (and slightly positive) in autumn compared to spring (0.2 ± 2.6 vs. -2.8 ± 2.2 µmol O_2_ g DW−^1^ h−^1^, respectively; Fig. 3A, Table S3), whereas NPP rates in the association showed no seasonal variation. Hourly respiration (R) rates were highest (more negative) in the sponge, followed by the seagrass-sponge association, and lowest in the seagrass (Fig. 3B, Table S2). Again, a seasonal effect was pronounced in the seagrass, with R rates in autumn 2-fold higher (more negative) than in spring (-3.8 ± 0.4 vs. -1.8 ± 0.3 µmol O_2_ g DW−^1^ h−^1^, respectively; Fig. 3B, Table S3), as well as in the sponge (-6.1 ± 1.1 vs. -3.6 ± 1.3 µmol O_2−_ g DW−^1^ h−^1^, respectively; Fig. 3B, Table S3). R rates in the association showed little seasonal variation (Fig. 3B). Daily rates of net community production (NCP) were highest in the seagrass treatment across both seasons, while the association showed intermediate values, and the sponge exhibited negative values (Fig. 3C, Table S2). Daily NCP was 54% lower in autumn compared to spring in the seagrass (87.2 ± 38.0 vs. 189.0 ± 37.7 µmol O_2−_ g DW−^1^ day−^1^, respectively; Fig. 3C, Table S3), while it remained stable across seasons in the sponge (-70.8 ± 28.8 vs. -76.0 ± 28.7 µmol O_2−_ g DW−^1^ day−^1^, respectively; Fig. 3C, Table S3) and in the association (73.6 ± 29.5 vs. 78.8 ± 19.6 µmol O_2−_ g DW−^1^ day−^1^, respectively; Fig. 3C, Table S3).

**Figure 3.**
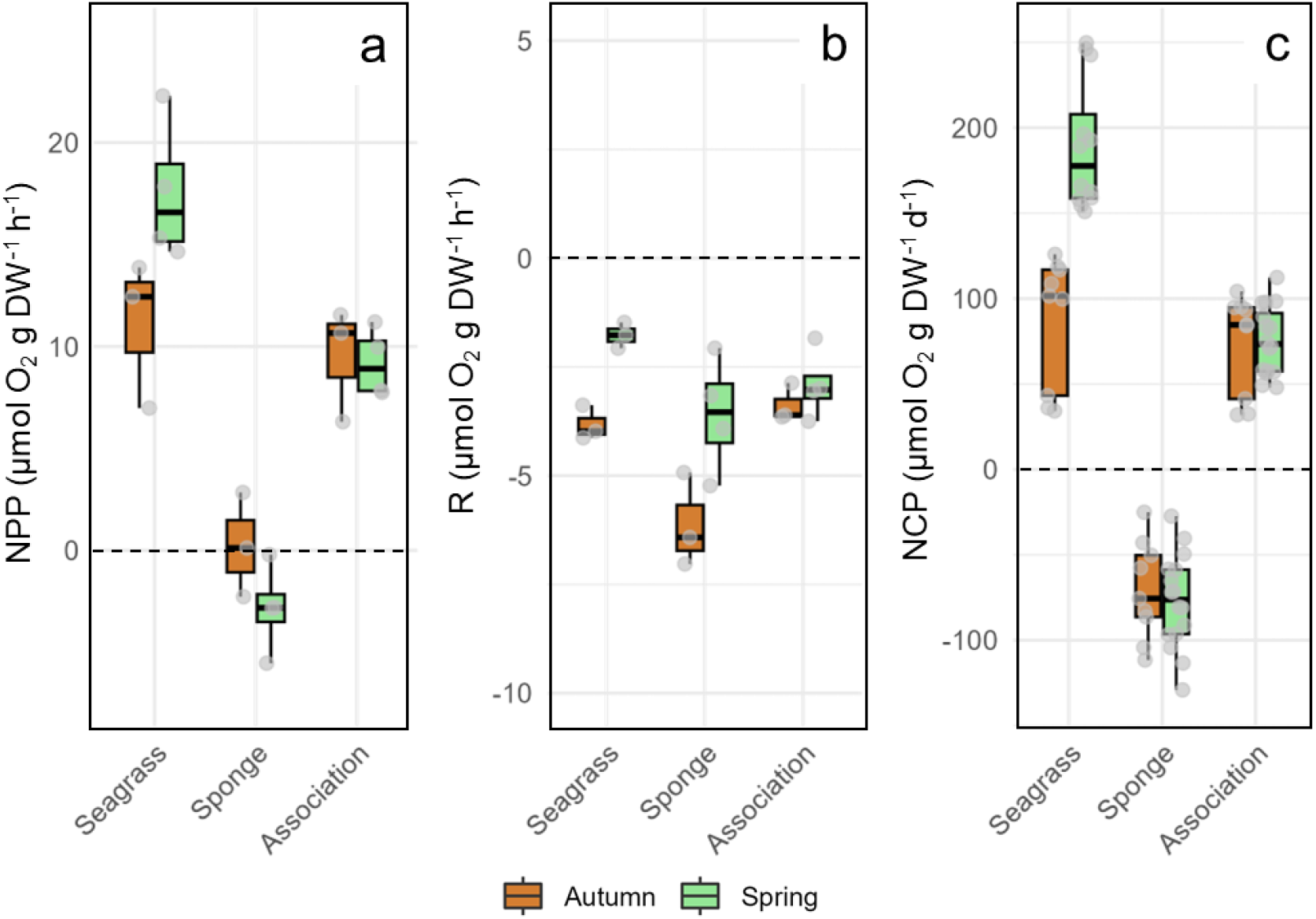
Seasonal variability in net primary production – NPP (a), dark respiration – R (b), and daily net community production – NCP (c) across the different *‘Community’* types (*P. oceanica, C. nucula*, and their association). Each panel displays the distribution of metabolic rates for two seasons (Autumn and Spring). Boxplots represent the interquartile range (IQR) with the median marked by a horizontal line. Whiskers extend to 1.5 × IQR, and outliers are shown as individual points (jittered for clarity). Positive values indicate oxygen production, while negative values reflect oxygen consumption. NPP and R are expressed in µmol O_2_ g DW−^1^ h−^1^, while NCP is expressed in µmol O_2_ g DW−^1^ d−^1^.

### Organic nutrient fluxes

Daily DOC fluxes (Fig. 4A, Table S4) showed the highest release rates in the seagrass treatment for both seasons, particularly in spring (autumn = 40.2 ± 7.9, spring = 63.1 ± 17.3 µmol C g DW−^1^ day−^1^, Table S5). In contrast, the sponge exhibited DOC release in autumn and uptake in spring (27.3 ± 8.6 µmol C g DW−^1^ day−^1^ and -28.4 ± 24.9 µmol C g DW−^1^ day−^1^, respectively, Table S5). The association also showed seasonality in DOC fluxes, with significant release in autumn (40.9 ± 18.2 µmol C g DW−^1^ day−^1^) and a shift to minimal net uptake in spring (-1.4 ± 20.5 µmol C g DW−^1^ day−^1^; Fig. 4A, Table S5). For the most part, daily DON fluxes (Fig. 4B, Table S4) showed release across treatments and seasons. In autumn, the highest DON release was in the sponge (7.6 ± 2.4 µmol N g DW−^1^ day−^1^), with intermediate values in the association and the lowest in seagrass (3.5 ± 1.2 µmol N g DW−^1^ day−^1^, Table S5). During spring, the seagrass exhibited the highest DON release (10.8 ± 3.2 µmol N g DW−^1^ day−^1^), with intermediate values in the sponge and the lowest in the association (0.8 ± 1.7 µmol N g DW^-1^ day^-1^; Fig. 4B, Table S5). Hourly DOC fluxes (Fig. S1, Table S4, S5) were higher in the light than in the dark in the seagrass and association *Community* types, with highest values for the seagrass in the light in spring. The sponge showed release in autumn but uptake in spring, with no clear differences between dark and light. Hourly DON fluxes were higher in the dark than in the light in autumn for all three *Community* types (Fig. S1, Table S4, S5), while they were higher in the light than in the dark in the seagrass and in the association in spring.

**Figure 4.**
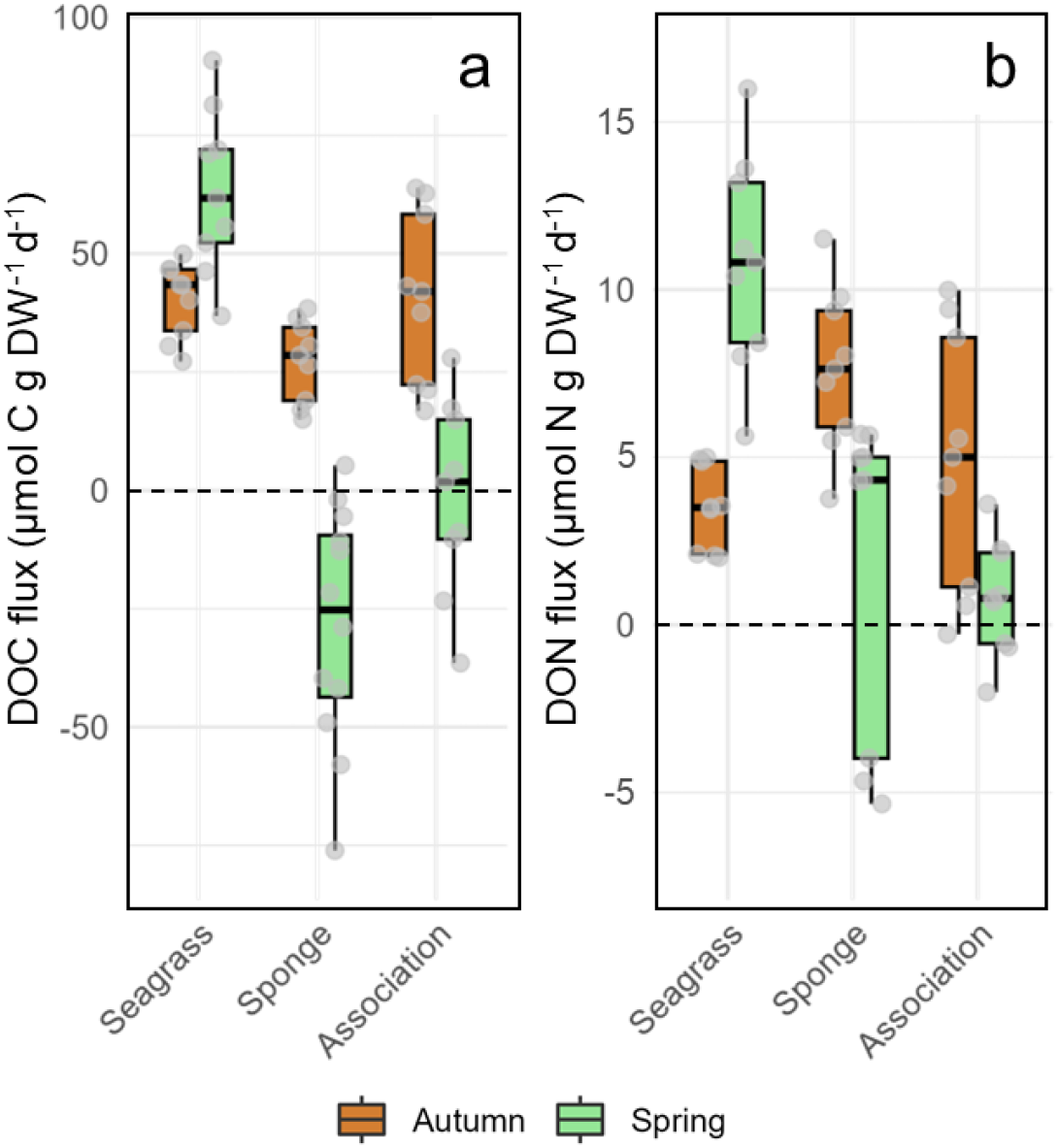
Seasonal variability in dissolved organic carbon – DOC (a) and dissolved organic nitrogen – DON (b) daily fluxes across the different *‘Community’* types (*P. oceanica, C. nucula*, and their association). Each panel shows the distribution of nutrient fluxes for two seasons (Autumn and Spring). Boxplots represent the interquartile range (IQR) with the median indicated by a horizontal line. Whiskers extend to 1.5 × IQR, with points outside this range plotted as individual data points (jittered for visibility). Negative values indicate nutrient uptake, and positive values indicate release. Flux rates are expressed in µmol g DW−^1^ d−^1^.

### Inorganic nutrient fluxes

Daily 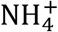 fluxes (Fig. 5A) showed constant uptake in the seagrass throughout the *Seasons* (autumn and spring = -0.3 ± 0.4 and -1.6 ± 0.5 µmol 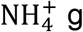 DW−^1^ day−^1^, respectively; Table S6, S7). Conversely, the sponge showed consistent 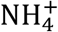 release (autumn and spring = 0.7 ± 0.6 and 2.4 ± 0.7 µmol 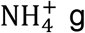 DW−^1^ day−^1^, respectively), while the association showed release in autumn (1.6 ± 0.6 µmol 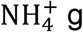 DW−^1^ day−^1^) and a flux close to net zero in spring (0.4 ± 0.9 µmol 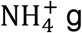 DW−^1^ day−^1^; Table S6, S7). The seagrass showed low but consistent 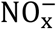 uptake (autumn = -0.2 ± 0.1, spring = -1.2 ± 0.3 µmol 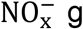 DW^-1^ day^-1^; Fig. 5B), while the sponge displayed strong and consistent 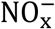 release (autumn = 13.6 ± 5.4, spring = 16.7 ± 5.5 µmol 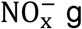 DW^-1^ day^-1^), with the association showing intermediate but positive fluxes in both seasons (autumn = 4.6 ± 1.9, spring = 6.5 ± 2.3 µmol 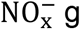 DW^-1^ day^-1^; Table S6, S7). In autumn, both the seagrass and the sponge showed 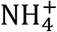 release in the light and 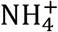 uptake in the dark (Fig. S2, Table S6,S7), while this pattern was not maintained in spring. Conversely, 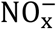 fluxes were highest in the sponge with higher release in the dark than in the daylight, across both autumn and spring (Fig. S2, Table S6, S7). 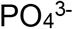 fluxes were low (close to detection limit) and variable, with no clear difference between daylight and dark but mainly uptake in the seagrass and release in the sponge and the association (Fig. S2, Table S6, S7).

**Figure 5.**
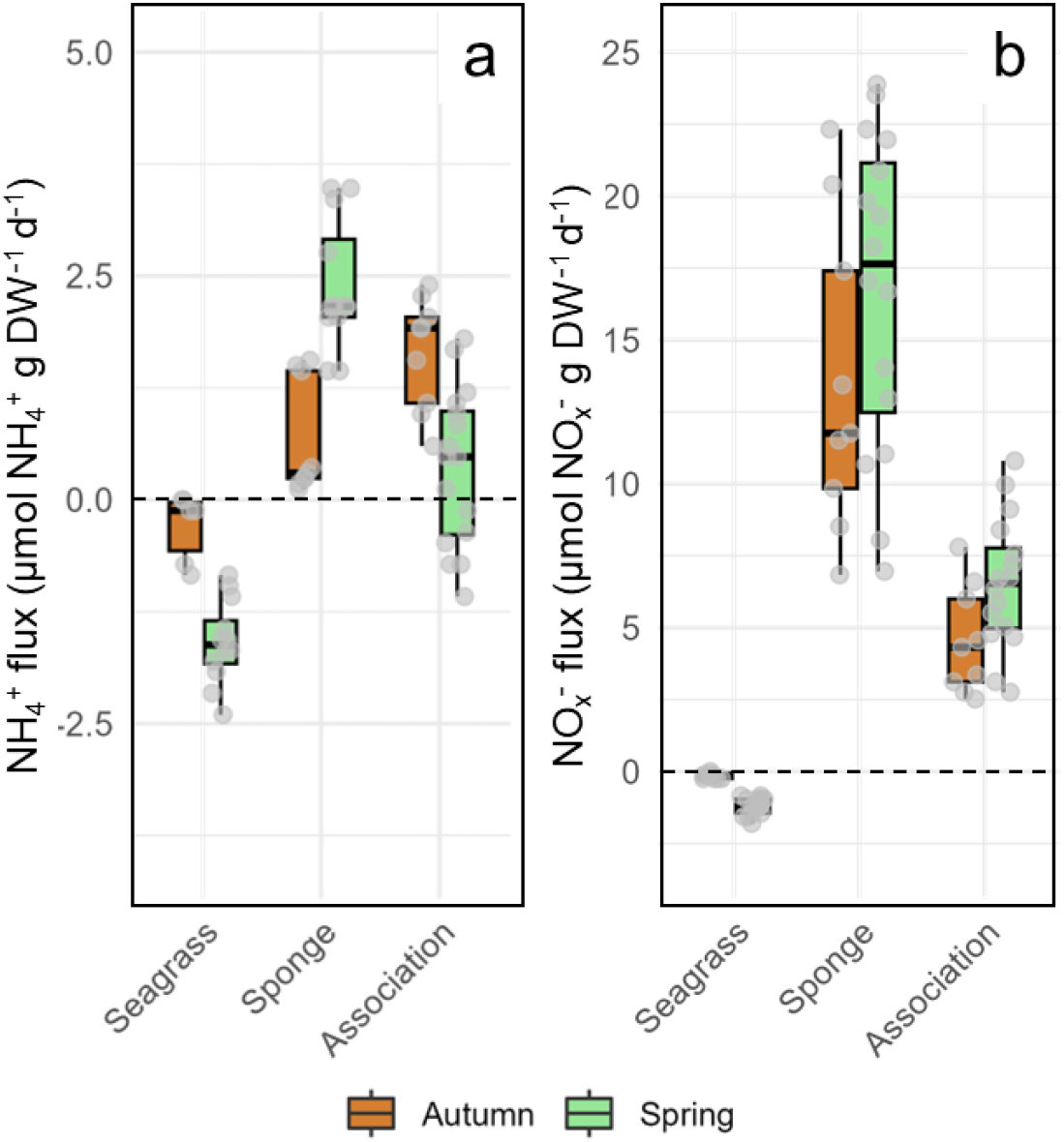
Seasonal variability in ammonium 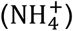, and nitrate + nitrite 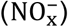 daily fluxes across the different *‘Community’* types (*P. oceanica, C. nucula*, and their association). Each panel shows the distribution of nutrient fluxes for two seasons (Autumn and Spring). Boxplots represent the interquartile range (IQR) with the median indicated by a horizontal line. Whiskers extend to 1.5 × IQR, with points outside this range plotted as individual data points (jittered for visibility). Negative values indicate nutrient uptake, and positive values indicate release. Flux rates are expressed in µmol g DW−^1^ d−^1^.

### Stable isotope analysis

δ^13^C of the seagrass ranged from -13.7 ± 1.3‰ when associated with the sponge to -13.9 ± 0.8‰ when non-associated, with no statistical differences between the two groups (n = 20; Fig. S3, Table S8). Concurrently, we detected an increase in δ^15^N of ca. 1‰ in *P. oceanica* living associated with the sponge, from 4.3 ± 1.0‰ to 5.3 ± 0.7‰ (Fig. S3, Table S8, S9). The sponge *C. nucula* showed similar δ^13^C and δ^15^N values for non-associated sponges (-19.2 ± 0.5‰ and 6.6 ± 0.7‰) vs sponges associated with the seagrass (-19.0 ± 0.5‰ and 6.7 ± 0.6‰, n=20; Fig. S3). Plant epiphytes had δ^13^C values similar to those of the sponge (-18.8 ± 2.4‰ and -19.6 ± 2.2‰, n=10, for associated vs non-associated plants, respectively) but showed an increase in their δ^15^N of ca. 1‰ when the plant was associated with the sponge (from 5.9 ± 1.0‰ to 7.3 ± 1.5‰, Fig. S3, Table S8, S9).

## DISCUSSION

This study provides new insights into an association between the seagrass *P. oceanica* and the sponge *C. nucula*, revealing key aspects of their interaction. We found that the association displays non-linear spatial dependence, with higher sponge abundance linked to intermediate seagrass cover. Further, we found evidence that the sponge benefited from DOC released by the seagrass in spring, while contributing substantial inorganic nitrogen, which may support seagrass productivity. These findings support the hypothesis of a facultative mutualism between *P. oceanica* and *C. nucula*, advance our understanding of the ecological dynamics within *P. oceanica* meadows, and highlight the importance of sponges in maintaining meadow stability and nutrient cycling.

### The sponge-seagrass association shows spatial dependence

We found evidence that the association between *C. nucula* and *P. oceanica* displays non-linear spatial dependence, with the maximum sponge cover occurring at intermediate seagrass cover (∼75%) and areas of both low and high seagrass cover corresponding to minimal sponge presence. This pattern suggests that at intermediate levels of seagrass cover, there is a favorable balance of available substrate and resource availability for both organisms at this site. These results align with SK Archer, EW Stoner and CA Layman [21], who reported that intermediate seagrass cover offers sufficient substrate for sponge colonization without significantly reducing water flow or light, which could otherwise impair sponge nutrition and/or seagrass photosynthesis. This is relevant because *C. nucula*, similarly to its congeneric species from the Caribbean and Australia, is a photophilic HMA sponge with a rich microbiome dominated by autotrophic cyanobacteria, which contribute to its energy production [25, 28, 29]. However, a previous report found that photoautotrophy could account for only a small fraction of the total daily carbon uptake in the Caribbean congeneric *C. caribensis* (ca. 7%), while DOC uptake contributed the most to the sponge diet (ca. 92%, [29]). At our study site, this may provide a competitive advantage for *C. nucula* to associate with a large primary producer, such as *P. oceanica*, which is known to release large amounts of DOM [30]. The asymmetric spatial dependence of the sponge and seagrass indicates neutrality for *P. oceanica* toward the presence of the sponge. However, this neutrality could be the result of a balance between positive and negative effects, rather than the absence of interaction, as KA Mathis and JL Bronstein [31] suggested. In particular, the seagrass may compete with the sponge for space, while at the same time may benefit from its efficient nutrient recycling capacity.

### Association with the sponge stabilizes meadow productivity

NPP measurements show that the sponge was near zero metabolic balance in autumn but shifted to net heterotrophy in spring. In contrast, the seagrass and the seagrass-sponge association remained autotrophic throughout both seasons. Our respirometry results are consistent with rates and seasonal dynamics described in previous studies [32-34], providing confidence that the data collected on these occasions are representative of broader seasonal processes, and confirming a shift for the plant from a highly productive growth phase in spring to a senescent phase in autumn. Sponges where symbiont photosynthesis exceeds holobiont respiration are termed “net phototrophic” and are estimated to have >50% of their daily respiratory needs met by their photosynthetic partners [35]. Although microbial diversity in *C. nucula* was not assessed in this study, our rates are to be attributed to cyanobacterial symbiont photosynthesis [28], which contributed significantly to sponge nutrition, providing approximately 52% of daily respiratory carbon demand in autumn while only a minor fraction in spring (Fig. 6). This underscores the importance of mixotrophy in this sponge, which, similar to what has been reported for a congeneric species in the Caribbean, may rely heavily on DOM uptake [29]. *P. oceanica* showed strong seasonality in productivity (NPP and NCP), but this variation was significantly less pronounced when *C. nucula* was present, indicating a buffering effect by the sponge due to its increased autotrophy in autumn. It is known that biodiversity can enhance productivity, resource use, and stability of seagrass ecosystems [36]. Similarly to land plants, where species interactions that present asynchrony in species fluctuations result in niche partitioning or facilitation and increase both productivity and temporal stability [37], meadows colonized by *C. nucula* may exhibit lower primary production relative to non-colonized meadows during productive seasons, but increased sponge activity during the plant senescence season may buffer the ecosystem against nutrient limitations, favoring nutrient recycling and promoting long-term stability.

**Figure 6.**
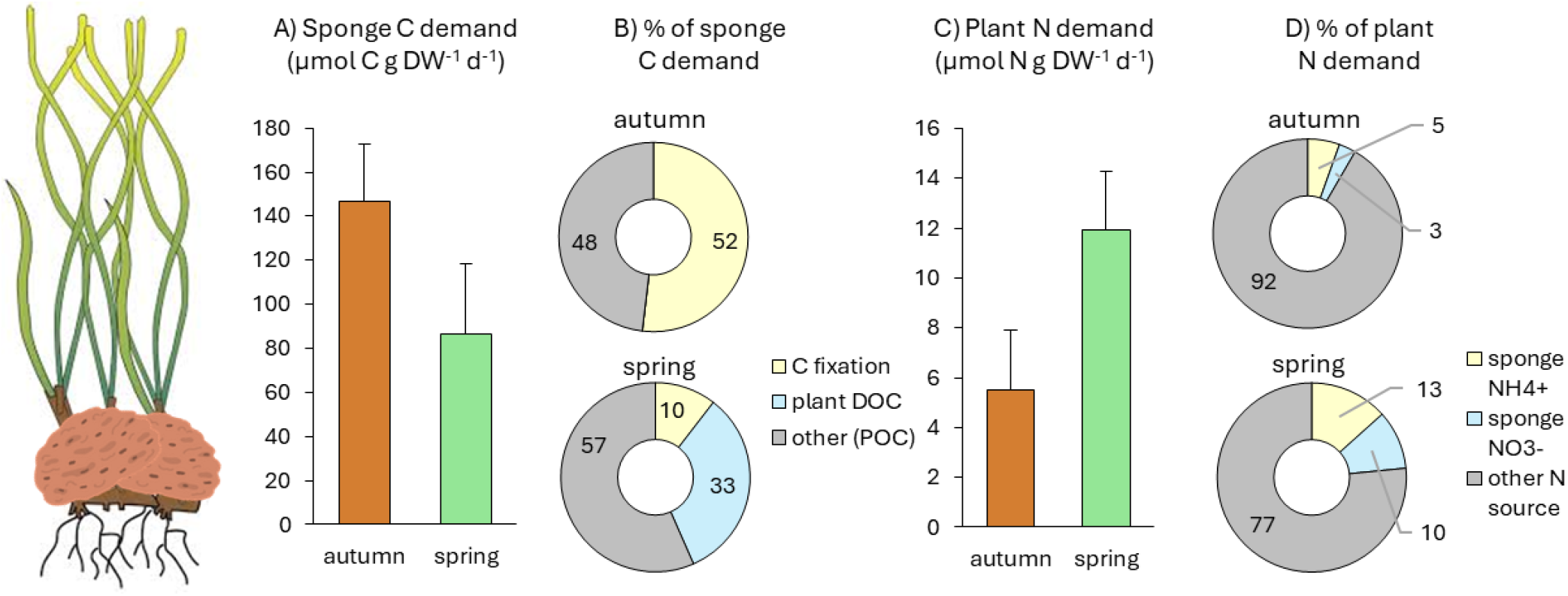
Photoautotrophic and heterotrophic nutrient recycling within the *P. oceanica*–*C. nucula* association. (A) Seasonal variation in the sponge daily respiratory carbon (C) demand (µmol C g DW−^1^ d−^1^), shown as mean ± SD. (B) Potential contribution (%) of heterotrophic dissolved organic carbon (DOC) and photoautotrophic C fixation to the sponge respiratory C demand, with the remaining fraction hypothesized to originate from particulate organic carbon (POC) via filter-feeding. (C) Seasonal variation in the plant daily nitrogen (N) demand estimated from net community production (NCP) and C:N ratios (µmol N g DW−^1^ d−^1^), shown as mean ± SD. (D) Potential contribution (%) of sponge ammonium 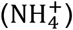 and nitrate+nitrite 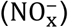 release to the plant total daily N demand. See the supplementary methods for details.

### Nutrient fluxes between seagrass and sponge underpin the association

*P. oceanica* contributed significant amounts of DOM (as DOC and DON) to its surrounding environment, particularly in spring, concomitant with the highest plant NPP rates. *P. oceanica* is known to enhance DOC fluxes relative to adjacent unvegetated sediments, as these plants produce nonstructural carbohydrates in excess [30, 38]. In particular, we estimate that the plant released approximately 46% and 33% of its NCP as DOC in autumn and spring, respectively. This is lower than the 71% estimate by C Barrón and CM Duarte [30], for a *P. oceanica* community in Mallorca Island (Spain), although their estimate also included contributions from allochthonous inputs.

DON was for the most part released by all community types across both seasons, albeit with high variability. The pattern mirrored that of DOC fluxes, with the highest DON release observed in seagrass during spring, coinciding with its growth season, and a tendency for DON uptake by the sponge and the association also in spring. DON fluxes in benthic organisms such as seagrasses and sponges have been rarely documented. The rates measured here are lower but comparable to those reported by S Liu, Z Jiang, C Zhou, Y Wu, I Arbi, J Zhang, X Huang and SM Trevathan-Tackett [39] for the tropical seagrasses *Thalassia hemprichii* and *Enhalus acoroides*, attributed to the leaching of nonstructural carbohydrates and other labile organic matter from seagrass leaves. In addition to seagrass leaching, epiphytes and sponges may also contribute DON to the surrounding environment. These sources of DON may have supported sponge heterotrophy in spring, as well as microbial processes such as ammonification [40] and nitrification [41], thereby facilitating nitrogen cycling within the studied system.

Concurrently, we detected net DOC uptake by *C. nucula* during spring, under both light and dark conditions, while the sponge released DOC in autumn. DOC uptake/release aligns with the sponge’s mixotrophic condition, shifting between autotrophy-dominated in autumn (DOC release) and heterotrophy-dominated (DOC uptake) in spring, as indicated by our measurements of NPP. If we assume linear DOC removal by the sponge in response to rising DOC concentrations in the environment [42] and similar DOC release rates by the plant when associated with the sponge compared to when it is not associated, we can estimate that the DOC released by the seagrass in spring may have covered approximately 33% of the sponge’s respiratory carbon demand (Fig. 6), a decrease from 92% in the congeneric low-light dwelling *C. caribensis* [29]. Given the difference in depth niches between the two species and the shallow depth at our study site, it is reasonable to expect that *C. nucula* relies more on photoautotrophy compared to its congeneric species from the Caribbean. This reliance on photoautotrophy was estimated to cover approximately 52% of its respiratory carbon needs in autumn and around 10% in spring (Fig. 6).

While the seagrass contributed DOC, which was taken up by the sponge in spring, the sponge excreted significant amounts of 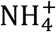 and 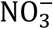 potentially benefiting plant growth. Indeed, we measured substantial uptake of both 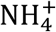 and 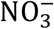 by the seagrass, particularly during spring, which coincides with its peak productivity season. HMA sponges are particularly known for contributing dissolved inorganic nutrients to their surroundings, primarily in the forms of 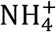,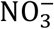, and 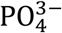 [14]. Notably, *C. nucula* exhibited substantial NOx− fluxes in both autumn and spring. These fluxes are likely the result of microbial nitrification within the sponge’s body. Numerous studies have demonstrated that sponges often associate with ammonium-oxidizing and nitrite-oxidizing microorganisms [43-45], and *C. nucula* at our study site is probably no exception. Specifically 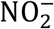 and 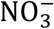 production rates in *C. nucula* aligned closely with those reported in previous studies from the Caribbean [46, 47], and showed that 98% of 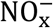 was released as 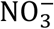. This pattern indicates the coexistence of ammonia-oxidation and nitrite-oxidation within the sponge holobiont. Further studies are needed to quantify these processes using stable isotope labeling. However, if we conservatively assume that the plant’s uptake remains constant when in association with the sponge compared to when it is not, we can estimate that the sponge contributed approximately 10% to the plant N demand in spring through 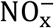 uptake and about 13% through 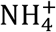 uptake (Fig. 6).

Our analysis of stable isotope data reveals that both the plant and its epiphytes exhibited higher δ^15^N values when associated with the sponge compared to when they were found alone. Significant isotopic fractionation occurs during microbial nitrification, resulting in the product 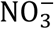 being depleted in δ^15^N while the residual 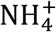 becomes enriched in δ^15^N [48]. In fact, the isotopic composition of 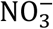 expelled from sponges *in situ* has lower δ^15^N values than 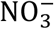 from the ambient water column due to nitrification [45]. Therefore, in our study, the increase in δ^15^N values in both the plant and its epiphytes when associated with the sponge may have resulted from the preferential incorporation of δ^15^N-enriched residual 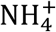 excreted by the sponge, which becomes enriched in ^15^N during microbial nitrification. Seagrasses often exhibit preferential uptake of 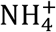 with 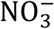 uptake rates representing only a small fraction of total nitrogen uptake [49, 50], also because 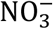 incorporation implies active transport with an associated energetic cost [16]. However, this pattern may not prevent the *P. oceanica* holobiont from benefiting also from the released 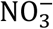. This seagrass species exhibits a complex nitrogen budget that involves both uptake and recycling processes [40, 41], which may enable *P. oceanica* to utilize 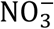 particularly in spring [51]. This capability may allow the plant to meet its nitrogen requirements even in the nutrient-poor conditions of the Mediterranean Sea.

## CONCLUSIONS

Our study highlights the ecological significance of the association between the seagrass *P. oceanica* and the sponge *C. nucula*, offering new evidence of spatial dependence and nutrient exchange between these organisms. The results suggest that intermediate seagrass cover promotes sponge colonization while ensuring favorable conditions for both organisms. The association is characterized by a dynamic seasonal balance between autotrophy and heterotrophy, with the sponge benefiting from seagrass-derived DOC and contributing inorganic nitrogen that likely enhances seagrass productivity in spring. This facultative mutualism stabilizes meadow productivity by buffering seasonal fluctuations and underscores the critical role of sponges in nutrient recycling within seagrass ecosystems. Future research should focus on tracing nutrient flows between the two species using stable isotope labeling, quantifying the contribution of microbial nitrification to the sponge’s nitrogen output, as well as assessing the stability of the association under environmental stressors. By deepening our understanding of these interactions, we can better predict how such associations may respond to global changes and contribute to the stability of seagrass ecosystems.

## METHODS

### Study area and benthic cover

Experiments were conducted in the “Schiacchetiello” inlet (40.7938 N, 14.0870 E, Southern Tyrrhenian Sea, Mediterranean), located in the municipality of Bacoli, Italy. Here, the seagrass *P. oceanica* grows at depths of 0-6 m, forming a patchy meadow. In many patches, the sponge *C. nucula* is found growing at the base of seagrass shoots, enveloping the rhizomes (Fig. 1A-D). We conducted video transects by snorkeling along the longest distance across the shallow (0.5 – 2 m) seagrass patches in November 2021. Video footage was processed using FFMPEG (https://ffmpeg.org/) to extract frames, from which 82 images were randomly selected from high quality images. We used these images to estimate the cover percentage of the primary benthic substrates within replicated 0.25 m^1^ quadrats, resulting in a total surveyed area of 20.5 m^1^. Inorganic and organic nutrient concentrations, as well as light intensity and temperature data, were collected upon each sampling occasion with methods as those reported in the following section to characterize environmental conditions at the study site where sampling took place for the following incubations.

### Oxygen and nutrient fluxes

*P. oceanica* shoots (hereafter “seagrass”), *C. nucula* bundles (“sponge”), and seagrass shoots hosting *C. nucula* (“association”) were collected from shallow beds (∼1.5 m depth) for *in situ* closed-chamber incubations, as in CA Pfister, U Cardini, A Mirasole, LM Montilla, I Veseli, J-P Gattuso and N Teixido [40]. These experiments were performed during midday hours on one sampling occasion in each of two seasons: autumn (November 2021) and spring (May 2022). We acknowledge that the lack of within-season replication limits our ability to assess intra-seasonal variability. However, the study was designed to capture distinct differences between two key periods in the plant’s life cycle: the growth season (spring) and the senescence season (autumn). These represent critical phases of biological activity, and the experiments were intended to provide a comparative snapshot of these distinct functional states. In autumn, incubations were conducted using 0.55 L cylindrical chambers (n = 3) for ∼3.75 hours; in spring, we used 1.1 L chambers (n = 4) for ∼6 hours to adjust for longer leaf length in this season and to maintain a similar biomass to volume ratio. Control chambers (n = 3) containing only seawater were included to measure fluxes from the water column community. Another full set of chambers were wrapped in three layers of black polyethylene to block light and incubated as “dark” chambers alongside the “light” chambers. The chambers were held in floating crates to maintain exposure to natural sunlight (for the “light” chambers) and seawater temperature while allowing for gentle wave action to prevent stratification. HOBO data loggers were used to monitor light intensity and temperature both within the chambers and externally, ensuring conditions resembled those found *in situ*. Discrete measurements with a LICOR light sensor allowed verifying that light chambers in autumn (when light intensity was lower, see Table 1) received light levels well above saturation irradiance (∼ 400 µmol quanta m^-2^ s^-1^). At the beginning and end of each incubation, O_2−_ concentrations inside the chambers were measured using a portable digital meter (WTW Multi 3430 Set K). After the incubations, 30 mL of seawater from each chamber was collected for dissolved organic carbon (DOC) and nitrogen (DON) analysis, and 20 mL was collected for inorganic nutrient measurements (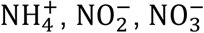 and 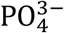). Samples were collected with acid-washed syringes and filtered immediately: DOC/DON samples were filtered using pre-combusted GF/F glass microfiber filters (pore size: 0.7 µm), acidified with 80 µL HCl (6 M), and refrigerated at 4°C until analysis. Inorganic nutrient samples were filtered through 0.22 µm PES membranes, frozen *in situ*, and stored at -20°C. DOC/DON was analyzed using a TOC-L Analyzer with TN unit (Shimadzu Corporation, Japan), while inorganic nutrients were measured with a continuous flow analyzer (Flowsys, SYSTEA SpA). Sponge and seagrass samples from each chamber were lyophilized for dry weight determination. Hourly flux rates for oxygen and nutrients were calculated as the change in analyte concentrations over incubation time corrected for the signal in the controls and the effective seawater volume in the chamber, standardized to the dry weight of the organisms and expressed in µmol analyte g DW−^1^ h−^1^.

### Stable isotope analyses

To examine potential signals of nutrient transfer in the sponge-seagrass association, in spring 2022 we collected samples of *P. oceanica* (leaves and epiphytes) and *C. nucula*, when growing associated vs non-associated. Epiphytes were gently scraped off seagrass leaves and stored in Eppendorf tubes. All samples were placed in acid-washed vials and lyophilized for 48 hours. Dried tissues were ground to a fine powder using a tissue lyser, then acid-fumed before being weighed into silver capsules for isotope analysis. Samples were analyzed using a Flash Elemental Analyzer (Thermo Scientific) equipped with a single reactor (1020°C), along with a MAT 253 Plus isotope ratio mass spectrometer (IRMS) interfaced with a Conflo IV system (Thermo Scientific, Bremen, Germany). The δ^13^C and δ^15^N values were normalized to Vienna Pee Dee Belemnite and atmospheric air, respectively, after correcting for blanks, ion source linearity, and standardizing against laboratory working standards and international reference materials (IAEA-600, IAEA-603). Precision was typically <0.1‰ for δ^13^C and 0.2‰ for δ^15^N. The molar C:N ratios (mol:mol) were calculated from C and N weights in the capsules (µg) and based on their respective molecular weights.

### Data analysis

Net primary production (NPP) and respiration (R) were determined based on hourly O_2−_ fluxes in light and dark incubations, respectively. To compute integrated rates of gross primary production (GPP = NPP + |R|), daily net community production (NCP = GPP × 12 -|R| × 24), and daily fluxes for each nutrient (Daily Flux = light flux × 12 + dark flux × 12), we generated analytical combinations of the observed values for light and dark fluxes, assuming equal duration of daylight or darkness. Each pair of independent values was combined using the respective formulas to compute the distribution of integrated rates (n = 9 for autumn, n = 16 for spring). This output provided a comprehensive distribution of the potential outcomes based on the input datasets. Daily rates were expressed in µmol analyte g DW−^1^ d−^1^. We examined potential asymmetries in the dependence between seagrass and sponge cover using the “qad” package [52] and modeled their relationship using a generalized additive model (GAM) with the “mgcv” package [53]. One-way permutation-based analyses of variance (PERMANOVA) using Euclidean distance were performed on each response variable [54] to test the effects of *Community* (seagrass, sponge, association) and *Season* (autumn, spring) on hourly and daily oxygen fluxes as well as daily nutrient fluxes, while separate PERMANOVAs assessed hourly nutrient fluxes with *Condition* (light, dark) as an additional factor. δ^13^C and δ^15^N values and C:N ratios were tested for differences among *Sample* types (*P. oceanica* leaves, *P. oceanica* epiphytes, *C. nucula*) and *Association* types (associated vs non-associated) using one-way PERMANOVA. All data analyses were performed in R [55].

## DATA AVAILABILITY

Data presented in this manuscript can be found at https://doi.org/10.5281/zenodo.14005723.

## ACKNOWLEDGMENTS

This research was supported by the Italian PRIN2022 project ENGAGE (grant n. 20223R4FJK) and PRIN-PNRR project BORIS (grant n. P2022R739J). We acknowledge the National Recovery and Resilience Plan (PNRR) by the European Union – Next Generation EU (National Biodiversity Future Centre – NBFC, project code CN00000033). We further acknowledge support from a Ph.D. fellowship to LMM funded by the Open University – SZN Ph.D. Program, a postdoctoral fellowship to GZ-H funded by the Agencia Nacional de Investigación y Desarrollo of Chile (ANID - Becas Chile), and a Ph.D. fellowship to JB cofounded by the Stazione Zoologica Anton Dohrn (SZN) and the University of Bremen. We thank Luigi Gallucci for his valuable support during fieldwork and logistical assistance, which greatly facilitated data collection and sample transport to the Lab.

