## Supplementary materials for "Reciprocal nutritional benefits in a sponge-seagrass association"

#### SUPPLEMENTARY METHODS

To evaluate the reciprocal nutritional benefits between the sponge and the seagrass, we estimated the sponge daily respiratory carbon (C) demand and the seagrass daily nitrogen (N) demand in Fig. 6. These estimates were derived from sponge respiration data and plant NCP, respectively, assuming a respiratory/photosynthetic quotient of 1, a 24 h cycle for sponge respiratory C demand, and a 12:12 h light/dark cycle and average plant C:N ratio of 16 for plant N demand (Fig. S3). To quantify the contribution of different C sources to the sponge respiratory C demand, we used the daily sponge DOC uptake and GPP rates to calculate the percentage of C demand met by heterotrophic DOC uptake and photoautotrophic C fixation, respectively. The remainder was attributed to particulate organic carbon (POC) uptake via filter-feeding. This approach was conservative because sponge DOC uptake was lower than seagrass DOC release, although it is possible that sponge DOC uptake (and its contribution to the sponge respiratory C demand) increased when in association with the seagrass. For the seagrass, we used daily plant $NH_4^+$ and $NO_x$ uptake rates to conservatively estimate the percentage of its total daily N demand potentially fulfilled by $NH_4^+$ and $NO_x^-$ released by the sponge. The remainder was attributed to other N sources. This approach was also conservative because seagrass uptake rates were lower than sponge release, although plant uptake rates (and their contribution to the plant N demand) may have increased when associated with the sponge.

### SUPPLEMENTARY FIGURES

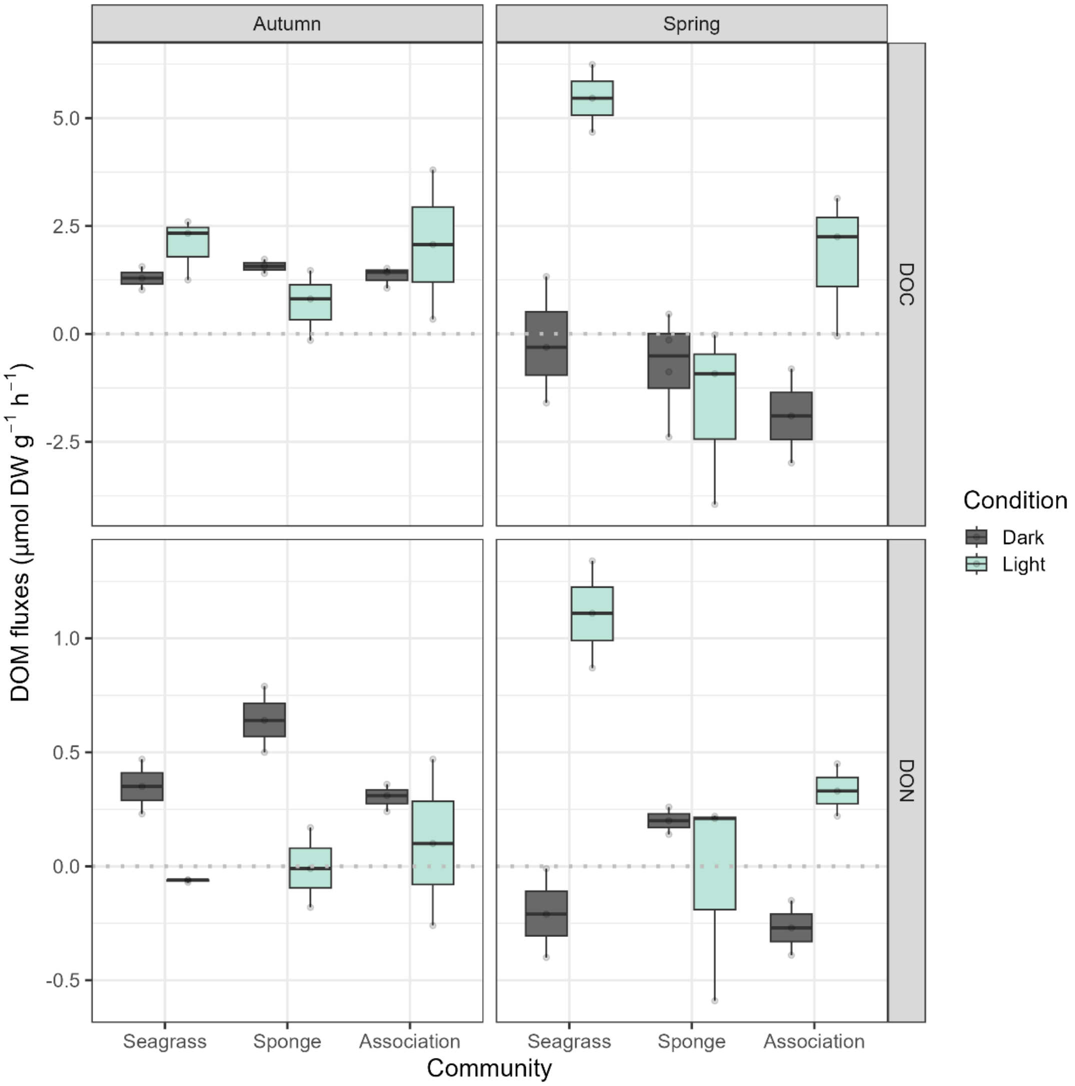


**Fig. S1.** Dissolved organic matter (DOM) fluxes expressed as µmol g $DW^{-1}$ $h^{-1}$ for dissolved organic carbon (DOC, top panels) and dissolved organic nitrogen (DON, bottom panels) across three communities—*Posidonia oceanica* (Seagrass), *Chondrilla nucula* (Sponge), and their Association. Fluxes are shown separately for two seasons: Autumn (left) and Spring (right). Each boxplot displays the fluxes under dark (gray) and light (teal) conditions. Positive values represent net release, while negative values indicate uptake. The horizontal dashed line marks the zero-flux threshold. Whiskers denote variability across replicates, and the central line in each box indicates the median flux.

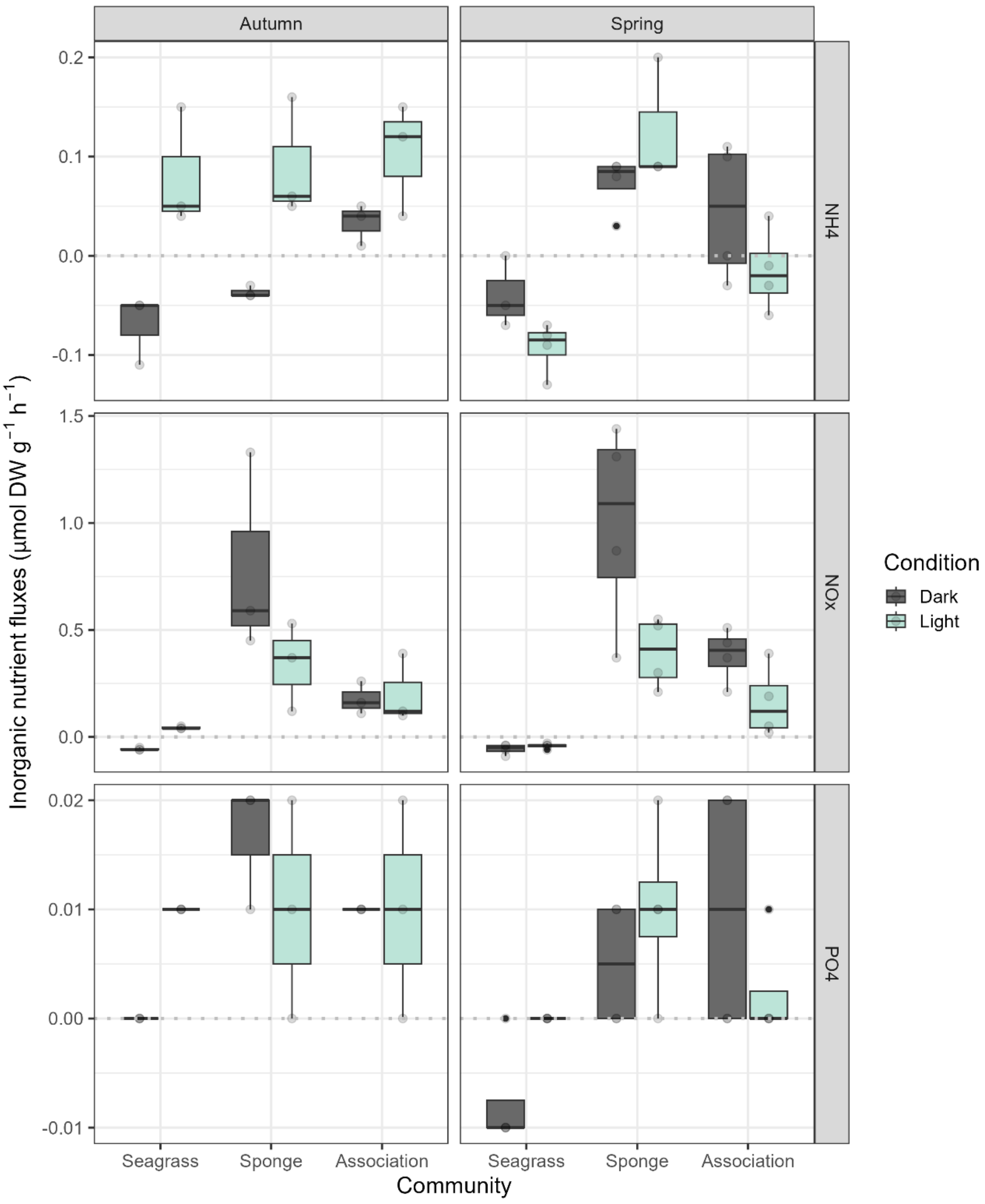


**Fig. S2.** Inorganic nutrient fluxes expressed as µmol g $DW^{-1}$ $h^{-1}$ for ammonium ($NH_4^+$, top panels), nitrate+nitrite ($NO_x^-$, middle panels), and phosphate ($PO_4^{3-}$, bottom panels) across three communities—*Posidonia oceanica* (Seagrass), *Chondrilla nucula* (Sponge), and their Association. Fluxes are presented separately for Autumn (left) and Spring (right) seasons under dark (gray) and light (teal) conditions. Positive values represent net release, while negative values indicate uptake. The horizontal dashed line marks the zero-flux threshold. Whiskers display variability across replicates, with the central line in each box representing the median flux.

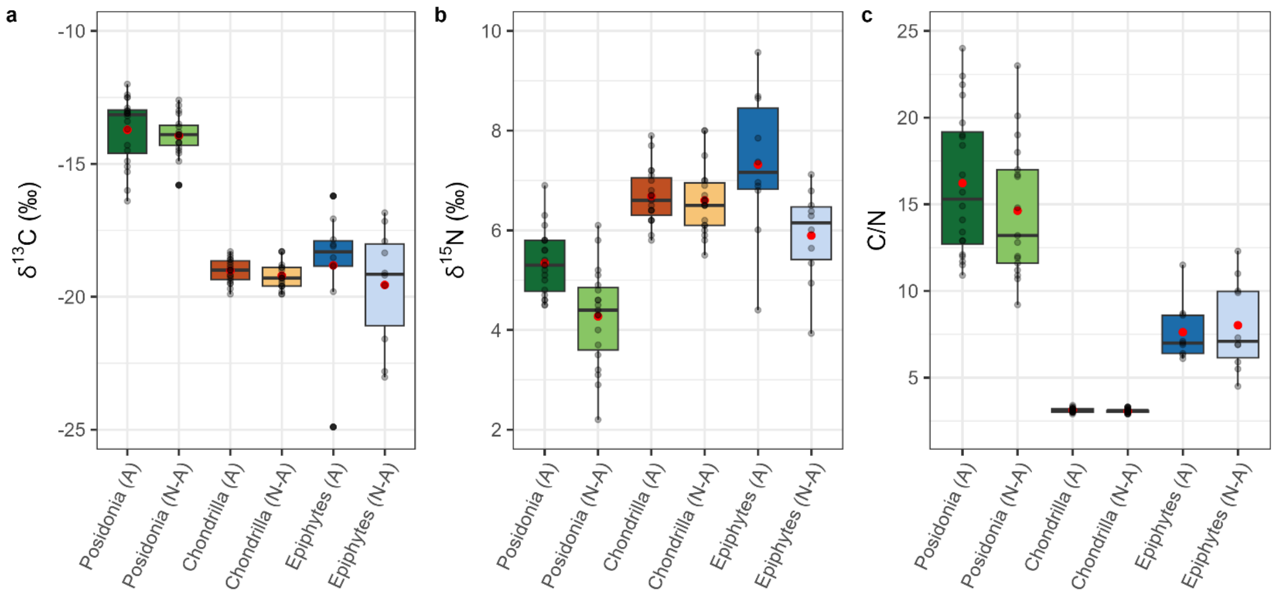


**Fig. S3.** Stable isotope composition and C/N ratios across communities in the association (A) and non-association (N-A) states. (a) $\delta^{13}C$ values (‰), (b) $\delta^{15}N$ values (‰), and (c) C/N ratios for *Posidonia oceanica*, *Chondrilla nucula*, and seagrass epiphytes. Each boxplot shows the distribution of values, with individual data points represented by black dots and the mean indicated by a red dot. Whiskers display the range of variability across replicates.

### SUPPLEMENTARY TABLES

**Table S1.** Coefficients of asymmetric dependency between the benthic cover of the seagrass *Posidonia oceanica* and the sponge *Chondrilla nucula*. The coefficient q(X,Y) indicates the strength of the dependency of organism Y on organism X. The asymmetry coefficient quantifies the imbalance between these dependencies, with a non-significant value suggesting that the interaction does not exhibit a strong directional asymmetry.

| **Coefficients** | **q** | **p-value** |
|---|---|---|
| q(*C. nucula*, *P. oceanica*) | 0.249 | **0.021** |
| q(*P. oceanica, C. nucula*) | 0.381 | **0.001** |
| asymmetry | -0.132 | 0.106 |

**Table S2.** Permutation-based analysis of variance (PERMANOVA) for net primary production (NPP, μmol $O_2$ g $DW^{-1}$ $h^{-1}$), respiration (R, μmol $O_2$ g $DW^{-1}$ $h^{-1}$), gross primary production (GPP, μmol $O_2$ g $DW^{-1}$ $h^{-1}$) and daily net community production (NCP, μmol $O_2$ g $DW^{-1}$ $d^{-1}$) across different *Community* types, *Seasons*, and their interaction. The table provides the degrees of freedom (Df), sum of squares (SS), proportion of variance explained ($R^2$), pseudo-F statistics, and associated p-values (P(>F)) for each source of variation. Bold p-values ($p < 0.05$) indicate which factors contribute to differences in the measured variables.

| Variable | Source of variation | Df | SS | $R^2$ | Pseudo-F | P (>F) |
|---|---|---|---|---|---|---|
| **Net primary production (NPP)** | Community | 2 | 963.6 | 0.83 | 63.48 | **0.001** |
| | Season | 1 | 5.3 | 0.00 | 0.70 | 0.389 |
| | Community:Season | 2 | 81.9 | 0.07 | 5.40 | **0.022** |
| | Residual | 15 | 113.9 | 0.10 | | |
| | Total | 20 | 1164.6 | 1.00 | | |
| **Respiration (R)** | Community | 2 | 13.6 | 0.33 | 9.20 | **0.001** |
| | Season | 1 | 13.7 | 0.33 | 18.51 | **0.001** |
| | Community:Season | 2 | 4.0 | 0.10 | 2.68 | 0.108 |
| | Residual | 14 | 10.4 | 0.25 | | |
| | Total | 19 | 41.7 | 1.00 | | |
| **Gross primary production (GPP)** | Community | 2 | 2588.8 | 0.79 | 208.26 | **0.001** |
| | Season | 1 | 12.7 | 0.00 | 2.05 | 0.159 |
| | Community:Season | 2 | 271.5 | 0.08 | 21.84 | **0.001** |
| | Residual | 65 | 404.0 | 0.12 | | |
| | Total | 70 | 3277.0 | 1.00 | | |
| **Net community production (NCP)** | Community | 2 | 592852 | 0.84 | 332.33 | **0.001** |
| | Season | 1 | 14804 | 0.02 | 16.60 | **0.003** |
| | Community:Season | 2 | 38754 | 0.06 | 21.72 | **0.001** |
| | Residual | 65 | 57978 | 0.08 | | |
| | Total | 70 | 704388 | 1.00 | | |

**Table S3**. Adjusted p-values from multilevel pairwise comparisons of net primary production (NPP), gross primary production (GPP) and net community production (NCP) between *Community* and *Season*. The comparisons are performed using Tukey's honest significant difference (HSD) test. Bold p-values indicate combinations that differ significantly ($p < 0.05$).

| **NPP** | Seagrass (Autumn) | Seagrass (Spring) | Sponge (Autumn) | Sponge (Spring) | Association (Autumn) | Association (Spring) |
|---|---|---|---|---|---|---|
| Seagrass (Autumn) | x | | | | | |
| Seagrass (Spring) | 0.067 | x | | | | |
| Sponge (Autumn) | 0.100 | **0.034** | x | | | |
| Sponge (Spring) | **0.029** | **0.032** | 0.136 | x | | |
| Association (Autumn) | 0.500 | **0.030** | 0.100 | **0.024** | x | |
| Association (Spring) | 0.356 | **0.025** | **0.031** | **0.030** | 0.855 | x |
| **GPP** | Seagrass (Autumn) | Seagrass (Spring) | Sponge (Autumn) | Sponge (Spring) | Association (Autumn) | Association (Spring) |
| Seagrass (Autumn) | x | | | | | |
| Seagrass (Spring) | **0.008** | x | | | | |
| Sponge (Autumn) | **0.001** | **0.001** | x | | | |
| Sponge (Spring) | **0.001** | **0.001** | **0.001** | x | | |
| Association (Autumn) | 0.150 | **0.001** | **0.002** | **0.001** | x | |
| Association (Spring) | **0.008** | **0.001** | **0.001** | **0.001** | 0.375 | x |
| **NCP** | Seagrass (Autumn) | Seagrass (Spring) | Sponge (Autumn) | Sponge (Spring) | Association (Autumn) | Association (Spring) |
| Seagrass (Autumn) | x | | | | | |
| Seagrass (Spring) | **0.001** | x | | | | |
| Sponge (Autumn) | **0.001** | **0.001** | x | | | |
| Sponge (Spring) | **0.001** | **0.001** | 0.602 | x | | |
| Association (Autumn) | 0.419 | **0.001** | **0.001** | **0.001** | x | |
| Association (Spring) | 0.320 | **0.001** | **0.001** | **0.001** | 0.858 | x |

**Table S4.** PERMANOVA for hourly (µmol g $DW^{-1}$ $h^{-1}$) and daily (µmol g $DW^{-1}$ $d^{-1}$) DOC and DON fluxes across different *Community* types, *Seasons*, *Condition* (light vs dark, when present) and their interaction terms. The table provides the degrees of freedom (Df), sum of squares (SS), proportion of variance explained ($R^2$), pseudo-F statistics, and associated p-values (P(>F)) for each source of variation. Bold p-values ($p < 0.05$) indicate which factors contribute to differences in the measured variables.

| Variable | Source of variation | Df | SS | $R^2$ | Pseudo-F | P (>F) |
|---|---|---|---|---|---|---|
| **Hourly DOC fluxes** | Community | 2 | 22.39 | 0.15 | 7.78 | **0.002** |
| | Season | 1 | 15.92 | 0.11 | 11.06 | **0.005** |
| | Condition | 1 | 16.66 | 0.11 | 11.58 | **0.003** |
| | Community:Season | 2 | 14.28 | 0.10 | 4.96 | **0.012** |
| | Community:Condition | 2 | 21.24 | 0.15 | 7.38 | **0.004** |
| | Season:Condition | 1 | 15.18 | 0.10 | 10.55 | **0.006** |
| | Community:Season:Condition | 2 | 7.30 | 0.05 | 2.54 | 0.106 |
| | Residual | 23 | 33.10 | 0.23 | | |
| | Total | 34 | 146.08 | 1.00 | | |
| **Hourly DON fluxes** | Community | 2 | 0.052 | 0.01 | 0.60 | 0.559 |
| | Season | 1 | 0.093 | 0.02 | 2.12 | 0.155 |
| | Condition | 1 | 0.000 | 0.00 | 0.00 | 0.998 |
| | Community:Season | 2 | 0.222 | 0.05 | 2.53 | 0.105 |
| | Community:Condition | 2 | 0.980 | 0.21 | 11.21 | **0.001** |
| | Season:Condition | 1 | 1.797 | 0.38 | 41.12 | **0.001** |
| | Community:Season:Condition | 2 | 0.533 | 0.11 | 6.10 | **0.014** |
| | Residual | 23 | 1.005 | 0.21 | | |
| | Total | 34 | 4.682 | 1.00 | | |
| **Daily DOC fluxes** | Community | 2 | 22408 | 0.36 | 36.83 | **0.001** |
| | Season | 1 | 11463 | 0.18 | 37.68 | **0.001** |
| | Community:Season | 2 | 14959 | 0.24 | 24.58 | **0.001** |
| | Residual | 45 | 13.691 | 0.22 | | |
| | Total | 50 | 62522 | 1.00 | | |
| **Daily DON fluxes** | Community | 2 | 77.45 | 0.09 | 4.09 | **0.028** |
| | Season | 1 | 44.62 | 0.05 | 4.71 | **0.032** |
| | Community:Season | 2 | 312.05 | 0.36 | 16.48 | **0.001** |
| | Residual | 45 | 426.04 | 0.50 | | |
| | Total | 50 | 860.15 | 1.00 | | |

**Table S5**. Adjusted p-values from multilevel pairwise comparisons of hourly DOC and DON fluxes between *Community* and *Condition*, and daily DOC and DON fluxes between *Community* and *Season*. The comparisons are performed using Tukey's honest significant difference (HSD) test. Bold p-values indicate combinations that differ significantly ($p < 0.05$).

| **Hourly DOC fluxes** | Seagrass (daylight) | Seagrass (dark) | Sponge (daylight) | Sponge (dark) | Association (daylight) | Association (dark) |
|---|---|---|---|---|---|---|
| Seagrass (daylight) | x | | | | | |
| Seagrass (dark) | **0.010** | x | | | | |
| Sponge (daylight) | **0.004** | 0.326 | x | | | |
| Sponge (dark) | **0.004** | 0.680 | 0.533 | x | | |
| Association (daylight) | 0.204 | 0.110 | **0.038** | 0.105 | x | |
| Association (dark) | **0.010** | 0.389 | 0.873 | 0.624 | 0.051 | x |
| **Hourly DON fluxes** | Seagrass (daylight) | Seagrass (dark) | Sponge (daylight) | Sponge (dark) | Association (daylight) | Association (dark) |
| Seagrass (daylight) | x | | | | | |
| Seagrass (dark) | 0.374 | x | | | | |
| Sponge (daylight) | 0.240 | 0.609 | x | | | |
| Sponge (dark) | 0.812 | 0.085 | **0.027** | x | | |
| Association (daylight) | 0.621 | 0.425 | 0.181 | 0.225 | x | |
| Association (dark) | 0.247 | 0.765 | 0.808 | 0.058 | 0.297 | x |
| **Daily DOC fluxes** | Seagrass (Autumn) | Seagrass (Spring) | Sponge (Autumn) | Sponge (Spring) | Association (Autumn) | Association (Spring) |
| Seagrass (Autumn) | x | | | | | |
| Seagrass (Spring) | **0.006** | x | | | | |
| Sponge (Autumn) | **0.006** | **0.002** | x | | | |
| Sponge (Spring) | **0.001** | **0.001** | **0.002** | x | | |
| Association (Autumn) | 0.924 | **0.049** | 0.056 | **0.001** | x | |
| Association (Spring) | **0.001** | **0.001** | **0.001** | **0.011** | **0.002** | x |
| **Daily DON fluxes** | Seagrass (Autumn) | Seagrass (Spring) | Sponge (Autumn) | Sponge (Spring) | Association (Autumn) | Association (Spring) |
| Seagrass (Autumn) | x | | | | | |
| Seagrass (Spring) | **0.001** | x | | | | |
| Sponge (Autumn) | **0.001** | 0.206 | x | | | |
| Sponge (Spring) | 0.273 | **0.006** | **0.004** | x | | |
| Association (Autumn) | 0.335 | **0.036** | 0.080 | 0.137 | x | |
| Association (Spring) | **0.004** | **0.001** | **0.001** | 0.559 | **0.014** | x |

**Table S6.** PERMANOVA for hourly (µmol g DW⁻¹ h⁻¹) and daily (µmol g DW⁻¹ d⁻¹) $NH_4^+$, $NO_x^-$, $PO_4^{3-}$ fluxes (µmol g DW⁻¹ h⁻¹) across different *Community* types, *Seasons*, *Condition* (light vs dark, when present) and their interaction terms. The table provides the degrees of freedom (Df), sum of squares (SS), proportion of variance explained ($R^2$), pseudo-F statistics, and associated p-values (P(>F)) for each source of variation. Bold p-values ($p < 0.05$) indicate which factors contribute to differences in the measured variables.

| Variable | Source of variation | Df | SS | $R^2$ | Pseudo-F | P (>F) |
|---|---|---|---|---|---|---|
| **Hourly $NH_4^+$ fluxes** | Community | 2 | 0.07 | 0.27 | 16.38 | **0.001** |
| | Season | 1 | 0.00 | 0.02 | 1.93 | 0.178 |
| | Condition | 1 | 0.01 | 0.05 | 6.44 | **0.026** |
| | Community:Season | 2 | 0.04 | 0.16 | 9.86 | **0.003** |
| | Community:Condition | 2 | 0.01 | 0.06 | 3.41 | **0.050** |
| | Season:Condition | 1 | 0.04 | 0.18 | 21.09 | **0.001** |
| | Community:Season:Condition | 2 | 0.01 | 0.03 | 1.58 | 0.233 |
| | Residual | 28 | 0.06 | 0.23 | | |
| | Total | 39 | 0.25 | 1.00 | | |
| **Hourly $NO_x^-$ fluxes** | Community | 2 | 3.27 | 0.56 | 31.06 | **0.001** |
| | Season | 1 | 0.06 | 0.01 | 1.22 | 0.298 |
| | Condition | 1 | 0.36 | 0.06 | 6.76 | **0.009** |
| | Community:Season | 2 | 0.08 | 0.01 | 0.72 | 0.509 |
| | Community:Condition | 2 | 0.52 | 0.09 | 4.91 | **0.013** |
| | Season:Condition | 1 | 0.05 | 0.01 | 0.96 | 0.347 |
| | Community:Season:Condition | 2 | 0.01 | 0.00 | 0.13 | 0.872 |
| | Residual | 28 | 1.47 | 0.25 | | |
| | Total | 39 | 5.82 | 1.00 | | |
| **Hourly $PO_4^{3-}$ fluxes** | Community | 2 | 0.001 | 0.29 | 9.24 | **0.002** |
| | Season | 1 | 0.000 | 0.08 | 4.93 | **0.040** |
| | Condition | 1 | 0.000 | 0.00 | 0.00 | 0.980 |
| | Community:Season | 2 | 0.000 | 0.04 | 1.13 | 0.328 |
| | Community:Condition | 2 | 0.000 | 0.05 | 1.62 | 0.204 |
| | Season:Condition | 1 | 0.000 | 0.00 | 0.06 | 0.797 |
| | Community:Season:Condition | 2 | 0.000 | 0.09 | 2.86 | 0.086 |
| | Residual | 28 | 0.001 | 0.45 | | |
| | Total | 39 | 0.002 | 1.00 | | |
| **Daily $NH_4^+$ fluxes** | Community | 2 | 79.80 | 0.58 | 86.21 | **0.001** |
| | Season | 1 | 0.89 | 0.01 | 1.92 | 0.189 |
| | Community:Season | 2 | 30.60 | 0.22 | 33.06 | **0.001** |
| | Residual | 58 | 26.84 | 0.19 | | |
| | Total | 63 | 138.13 | 1.00 | | |
| **Daily $NO_x^-$ fluxes** | Community | 2 | 2763.4 | 0.78 | 121.04 | **0.001** |
| | Season | 1 | 50.6 | 0.01 | 4.43 | **0.048** |
| | Community:Season | 2 | 53.8 | 0.02 | 2.35 | 0.116 |
| | Residual | 58 | 662.1 | 0.19 | | |
| | Total | 63 | 3529.8 | 1.00 | | |
| **Daily $PO_4^{3-}$ fluxes** | Community | 2 | 0.77 | 0.48 | 37.66 | **0.001** |
| | Season | 1 | 0.23 | 0.14 | 21.96 | **0.001** |
| | Community:Season | 2 | 0.03 | 0.02 | 1.59 | 0.186 |
| | Residual | 58 | 0.60 | 0.37 | | |
| | Total | 63 | 1.63 | 1.00 | | |

**Table S7**. Adjusted p-values from multilevel pairwise comparisons of hourly $NH_4^+$ and $NO_x^-$ fluxes between *Community* and *Condition*, and daily $NH_4^+$ fluxes between *Community* and *Season*. The comparisons are performed using Tukey's honest significant difference (HSD) test. Bold p-values indicate combinations that differ significantly ($p < 0.05$).

| **Hourly $NH_4^+$ fluxes** | Seagrass (daylight) | Seagrass (dark) | Sponge (daylight) | Sponge (dark) | Association (daylight) | Association (dark) |
|---|---|---|---|---|---|---|
| Seagrass (daylight) | x | | | | | |
| Seagrass (dark) | 0.427 | x | | | | |
| Sponge (daylight) | **0.023** | **0.002** | x | | | |
| Sponge (dark) | 0.359 | **0.019** | **0.035** | x | | |
| Association (daylight) | 0.285 | **0.018** | 0.110 | 0.802 | x | |
| Association (dark) | 0.197 | **0.005** | 0.059 | 0.628 | 0.886 | x |
| **Hourly $NO_x^-$ fluxes** | Seagrass (daylight) | Seagrass (dark) | Sponge (daylight) | Sponge (dark) | Association (daylight) | Association (dark) |
| Seagrass (daylight) | x | | | | | |
| Seagrass (dark) | 0.051 | x | | | | |
| Sponge (daylight) | **0.002** | **0.005** | x | | | |
| Sponge (dark) | **0.001** | **0.004** | **0.037** | x | | |
| Association (daylight) | **0.005** | **0.004** | **0.038** | **0.002** | x | |
| Association (dark) | **0.002** | **0.001** | 0.228 | **0.002** | 0.192 | x |
| **Daily $NH_4^+$ fluxes** | Seagrass (Autumn) | Seagrass (Spring) | Sponge (Autumn) | Sponge (Spring) | Association (Autumn) | Association (Spring) |
| Seagrass (Autumn) | x | | | | | |
| Seagrass (Spring) | **0.001** | x | | | | |
| Sponge (Autumn) | **0.002** | **0.001** | x | | | |
| Sponge (Spring) | **0.002** | **0.001** | **0.001** | x | | |
| Association (Autumn) | **0.001** | **0.001** | **0.009** | **0.027** | x | |
| Association (Spring) | 0.105 | **0.001** | 0.368 | **0.001** | **0.005** | x |

**Table S8.** PERMANOVA for δ$^{13}$C and δ$^{15}$N values and C:N ratios across different *Sample* types and *Association* types. The table provides the degrees of freedom (Df), sum of squares (SS), proportion of variance explained ($R^2$), pseudo-F statistics, and associated p-values (P(>F)) for each source of variation. Bold p-values ($p < 0.05$) indicate which factors contribute to differences in the measured variables.

| Variable | Source of variation | Df | SS | $R^2$ | Pseudo-F | P (>F) |
|---|---|---|---|---|---|---|
| | Sample | 2 | 658.12 | 0.82 | 208.75 | **0.001** |
| | Association | 1 | 2.35 | 0.00 | 1.49 | 0.236 |
| **δ$^{13}$C values** | Sample:Association | 2 | 1.15 | 0.00 | 0.37 | 0.700 |
| | Residual | 91 | 143.45 | 0.18 | | |
| | Total | 96 | 805.07 | 1.00 | | |
| | Sample | 2 | 75.99 | 0.46 | 50.64 | **0.001** |
| | Association | 1 | 13.83 | 0.08 | 18.44 | **0.001** |
| **δ$^{15}$N values** | Sample:Association | 2 | 7.56 | 0.05 | 5.04 | **0.005** |
| | Residual | 91 | 68.28 | 0.41 | | |
| | Total | 96 | 165.66 | 1.00 | | |
| | Sample | 2 | 3241.8 | 0.71 | 112.31 | **0.001** |
| | Association | 1 | 6.7 | 0.00 | 0.46 | 0.509 |
| **C:N ratios** | Sample:Association | 2 | 9.4 | 0.00 | 0.32 | 0.692 |
| | Residual | 91 | 1313.4 | 0.29 | | |
| | Total | 96 | 4571.3 | 1.00 | | |

**Table S9**. Adjusted p-values from multilevel pairwise comparisons of $\delta^{15}N$ values between *Sample* types and *Association* types. The comparisons are performed using Tukey's honest significant difference (HSD) test. Bold p-values indicate combinations that differ significantly ($p < 0.05$).

| **$\delta^{15}N$ values** | *P. oceanica* leaves (associated) | *P. oceanica* leaves (not associated) | *C. nucula* (associated) | *C. nucula* (not associated) | *P. oceanica* epiphytes (associated) | *P. oceanica* epiphytes (not associated) |
|---|---|---|---|---|---|---|
| *P. oceanica* leaves (associated) | x | | | | | |
| *P. oceanica* leaves (not associated) | **0.001** | x | | | | |
| *C. nucula* (associated) | **0.001** | **0.001** | x | | | |
| *C. nucula* (not associated) | **0.001** | **0.001** | 0.734 | x | | |
| *P. oceanica* epiphytes (associated) | **0.001** | **0.001** | 0.106 | 0.104 | x | |
| *P. oceanica* epiphytes (not associated) | 0.064 | **0.001** | **0.011** | **0.029** | **0.027** | x |
